# Hypoxia impacts small intestinal organoid stemness and differentiation

**DOI:** 10.1101/2023.12.30.573689

**Authors:** Xi Lan, Ping Qiu, Chunfeng Mou

## Abstract

Comprised of several cell types functioning differently, the small intestinal epithelial cells perform their functions and synergistically maintain homeostasis in the first part of the mammalian intestine. Due to the uneven distribution of the vessel, the oxygen level exhibits a gradient decreasing pattern in the normal intestine and becomes aberrant in some intestinal diseases. In this work, we find certain levels of hypoxia simulated by cobaltous chloride (CoCl_2_) caused an increase in the secretive cell types and a decrease in the absorptive cell types *in vitro* cultured mouse small intestinal organoids. Importantly, the intestinal stem cell amount is impacted which leads to attenuated epithelial regeneration. Our study highlights the cell-type-specific alterations under the hypoxia insult, which gives possible therapeutic hints for hypoxia-relevant gastrointestinal diseases.

## Introduction

The human small intestinal epithelium is the fastest self-renewing tissue, which renews every 3 to 5 days, with the structure of bending cells forming the crypts and villi (1). Intestinal stem cells (ISCs) reside at the crypt base and differentiate into the transit-amplifying (TA) compartment which divides 4-5 times within 12 hours (2). After that, the TA cells move upwards and differentiate into multiple cell types including enterocytes, goblet cells and enteroendocrine cells. During normal homeostasis or some intestinal injuries, ISCs can undergo self-renewal and proliferate to replenish cells that are lost to shedding (3, 4). The self-renewal and lineage differentiation into mature enterocytes, goblet cells and enteroendocrine cells are under the control of Wnt/β-catenin, Notch and bone morphogenetic protein (BMP) pathway signalings (5-7). Among them, Wnt/β-catenin signaling is of critical significance, and its downstream regulated gene Leucine-rich repeat-containing G-protein coupled receptor 5 (Lgr5) is identified as the marker for proliferative ISCs (8, 9).

Mutations that activate the Wnt/β-catenin pathway can result in human colorectal cancer (10, 11). Studies suggested Wnt as an essential growth factor for the maintenance of intestinal proliferation and homeostasis (12, 13), and ectopic Wnt signaling has been implicated in the initiation and progression of colorectal cancers (14, 15).

In a healthy human intestine, due to the counter-current blood and the existence of a large number of gut bacteria, the oxygen pressure (pO2) level is lower than the normal oxygen pressure (160 mmHg), therefore, it’s described as”physiologic hypoxia” (16). The local pO2 level decreases along the crypt-villus axis and the oxygen gradient becomes the lowest at the villus tip (17). This unique gradient physiologic oxygen pressure results in the stabilization of the transcription factor hypoxia-inducible factor (HIF) and the activation of hypoxia-relative signaling, which plays multiple roles in metabolism, mitochondrial autophagy and inflammation (18). However, in pathological conditions, profound hypoxic or anoxic are usually found in mucosal lesions (19). Hypoxia is also a feature of inflammatory bowel disease (IBD), and it affects inflammation in some intestinal diseases (16, 20). Studies have demonstrated the protective role of HIF on some intestinal diseases such as IBD and necrotizing enterocolitis (NEC) upon *in vivo* hypoxia injury (21-23), and the effect of hypoxia on intestinal cell response in intestinal cancer cells or upon radiotherapy and chemotherapy *in vitro* (24-26). In both scenarios, the intestinal stem cells (ISCs) have been shown to react by promoting intestinal recovery from injuries (27, 28). However, the mechanism of how ectopic hypoxia affects ISC function and how it impacts intestinal cell lineage differentiation is less clarified. Mazumdar et al. mentioned that HIF-1α regulates Wnt/β-catenin signaling in hypoxic embryonic stem (ES) cells and neural stem cells (29). Rohwer et al. demonstrated the role of HIF-1α in intestinal epithelial cells (IECs) during intestinal tumorgenesis, controlling Wnt/β-catenin activity and regulating metabolic reprogramming and inflammation (30). However, it remains to be elucidated how hypoxia impacts different cell types in intestine with or without affecting Wnt signaling.

Intestinal organoids isolated from adult stem cells have been established as an ingenious craft to study the human intestine (8). It’s a 3D model of intestinal tissue embedded in 3D matrices such as Matrigel and can be passaged and cultured for a long time. The organoids display the characteristic villi-crypt organization by heterogeneous epithelial cell monolayer, allowing it to recapitulate many features of normal gut epithelium with IECs comprising all types of epithelial cells (31). In addition, the organoid system is applied to estimate IEC functions *in vivo* such as barrier function and stem cell function (32, 33). Compared with the *in vivo* mouse model and 2D cell lines (26, 34) to study the influence of hypoxia on intestinal functions or tumorigenesis, intestinal organoids can be easily established from different staging of endoscopic biopsies in patients, making it a better model that can closely recapitulate the hypoxic intestine and a faster model to come up with therapeutic regimens.

Hypoxia exists in intestinal diseases such as IBD and NEC, wherein the mechanism by which hypoxia impacts ISC stemness and differentiation remains poorly understood. In the present study, we examined obvious changes of different cell lineages upon hypoxia treatment to intestinal organoids, as well as the upstream transcription factors regulating the ISC stemness. Our study further confirms organoids as the model to study intestinal diseases that are concurrent with hypoxia. This cobaltous chloride (CoCl_2_)-caused hypoxia-treated organoid model makes it a good method to study HIF-1α regulated signaling in intestine, helping the translation of basic science studies into IBD treatment.

## Result

### 1. Hypoxia treatment affects intestinal organoid stem cell renewal function

It’s known that the ability of IECs to form organoids and organoid regeneration after being passaged depends mainly on ISC function (4, 8). To investigate the effect of hypoxia exposure on intestine organoid stem cell regeneration capacity, we passaged the ISC-containing organoid as an assay of intestinal regenerative potential (8). Increasing dosages of CoCl_2_ (0, 10, 50, 100, 200 µM) were used to induce cellular hypoxia (35, 36), and did not impair the budding and morphology of organoids (Fig. 1A). The protein level of HIF-1α was upregulated 24 hours after hypoxia exposure (Fig. 1B). The organoid regeneration, which indicates stem cell function and generated from single intestinal organoid cells engendered fewer progenies in a gradient decreasing manner, as showed in the clonogenicity assay (Fig 1C).

**Figure 1.**
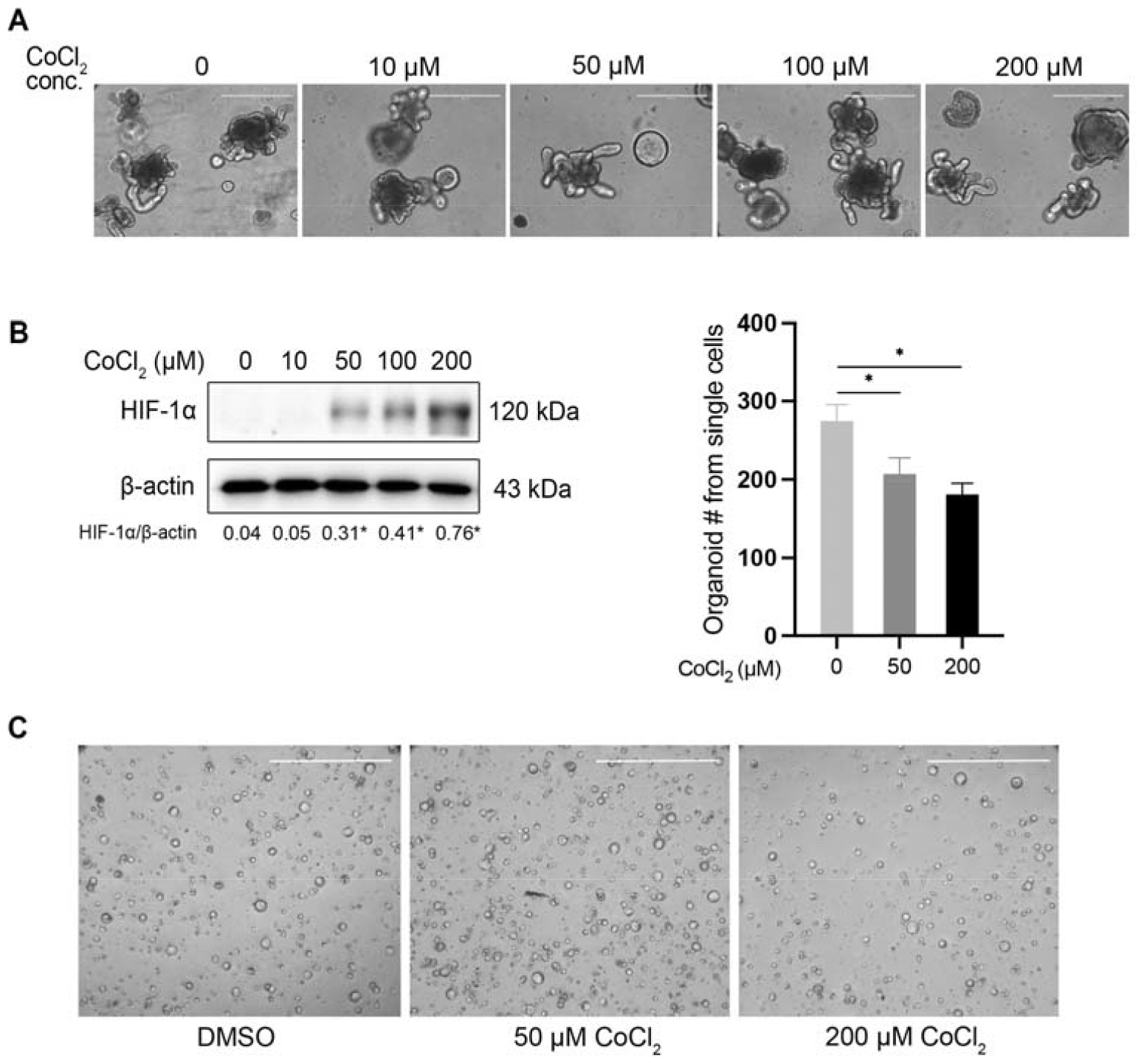
Hypoxia treatment affects intestinal organoid stem cell renewal function **(A)** Representative picture of organoids being treated with CoCl_2_ dissolved in DMSO at concentrations of 0, 10, 50, 100, and 200 µM. DMSO was used as the dissolvent and control treatment group. Scale bar, 400 µm. **(B)** Left, western blotting detection of HIF-1α and the number below shows the quantification of the band intensity by Image J. ^*^*P* < 0.05. Right, quantification of clonogenicity assay in panel C. **(C)** Clonogenicity of wild-type and hypoxia-treated organoids, 24 hrs post being seeded from single cells. In the control and CoCl_2_ treated group, the initial number of the single organoids is seeded at the same number. Scale bar, 1000 µm.

We further explored whether the reduction of the organoid regeneration function was preceded by the reduction of stem cell trait. Indeed, the mRNA levels of Lgr5 which mark the intestinal stem cells, were also down-regulated (Fig. 2A). Therefore, these data suggest that hypoxia exposure in intestine organoids impairs ISC regeneration by reducing ISC numbers and organoid-forming capacity.

**Figure 2.**
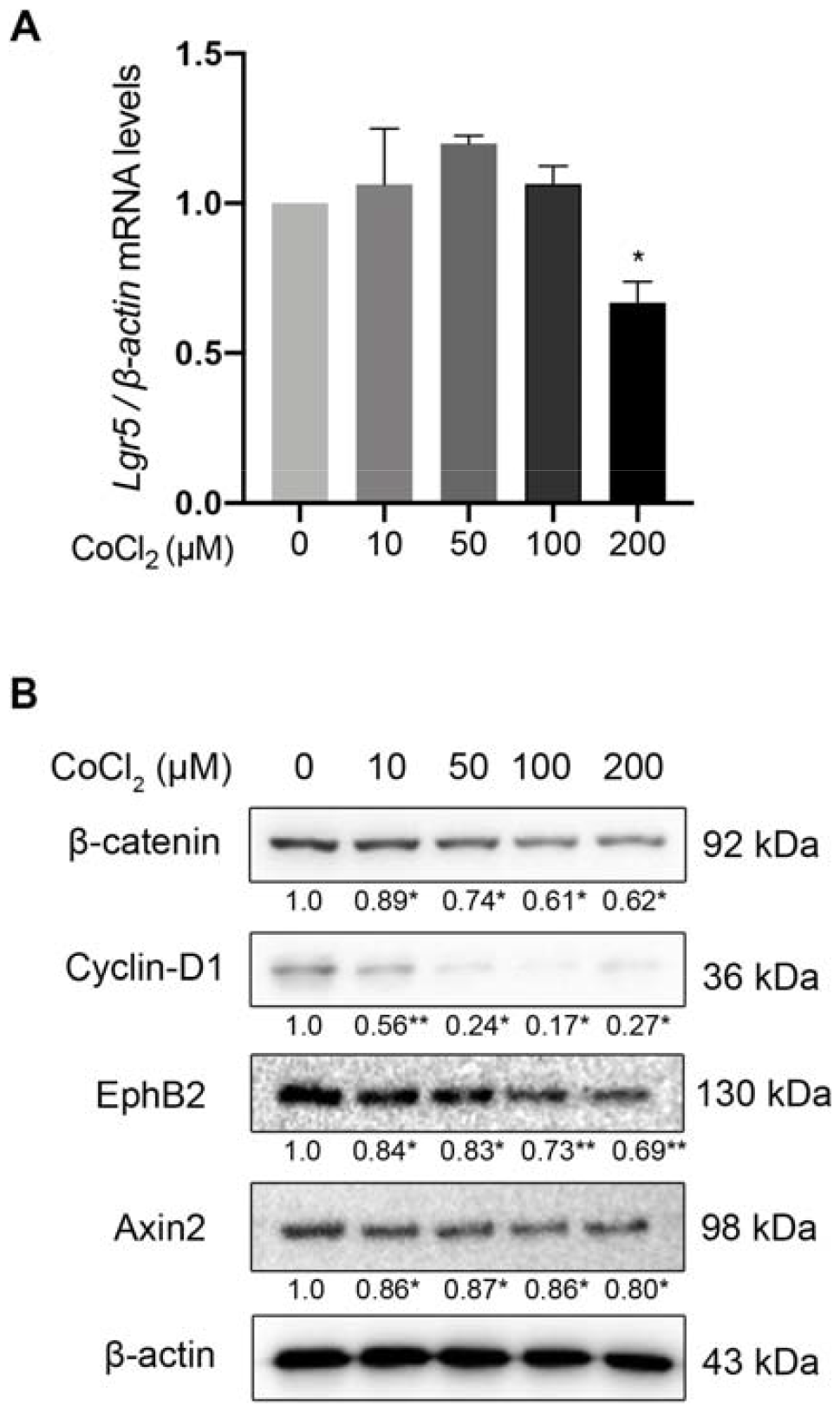
Wnt signaling was down-regulated upon hypoxia treatment in intestinal organoids **(A)** RT-qPCR detection of mRNA level of Lgr5 in organoids treated by gradient concentrations of CoCl_2_ for 24 h. ^*^*P* < 0.05. **(B)** Western blotting detection of Wnt targets (β-catenin, Cyclin-D1, Ephb2, and Axin2) in hypoxia-exposed organoids after increasing concentrations of CoCl_2_ treatment. ^*^*P* < 0.05, ^**^*P* < 0.01.

### 2. Wnt signaling was down-regulated upon hypoxia treatment in intestinal organoids

Wnt signals play a crucial role in the regulation of intestinal stem cell proliferation (11, 37, 38). As the key stem cell regulator in intestine stem cells, Lgr5 is controlled by Wnt/β-catenin signaling (4, 39) and exhibits a reduced mRNA level after 200 µM CoCl_2_ treated hypoxia treatment (Fig. 2A). To further clarify the effect of hypoxia on Wnt signaling in intestinal organoid, multiple Wnt target genes were tested in hypoxia-exposed intestine organoids. As the hypoxic condition gets severe, the protein level of β-catenin was down-regulated in a dosage-dependent manner (Fig 2B). Also, the β-catenin target Cyclin-D1, which is the gene involved in cell survival (40), exhibited significant protein level reduction. Expression of another Wnt target, the receptor tyrosine kinase EphB2, which reduces from the crypt base to the differentiated cells (41) and is highly enriched in ISCs (42), also shows a reduced protein level after 24 hrs hypoxia treatment. Consistent with this, protein expression of another Wnt/β-catenin target gene Axin2 (43) was also decreased. These demonstrate that hypoxia treatment dampens the activity of Wnt/β-catenin signaling in intestinal organoids.

### 3. Hypoxia treatment affects intestine differentiation and increases secretory cell types

Pathological conditions such as inflammation insults are usually accompanied by cellular hypoxia, further disrupting intestinal epithelial barrier function and metabolism (18). The barrier function of intestinal epithelium is largely composed of enterocytes to maintain intact epithelium construction, and goblet cell secreted mucus helps to form the healthy mucosa layer, making the epithelium devoid of bacteria (44).

Then we deciphered whether hypoxia drives different intestine cell lineage cell differentiation, and if it impairs certain cell types executing their functions. As shown in the mRNA levels, hypoxia exposure led to the increase of Lysozyme (Lyz), Chromogranin A (ChrA) and Mucin 2 (Muc2), which mark Paneth cell, enteroendocrine cell and goblet cell, respectively (Fig. 3A). At the same time, the protein expressions showed by the immunofluorescence (IFC) signal indicated a greater number of Lysozyme positive Paneth cells, Chromogranin A positive enteroendocrine cells, and Mucin2 positive goblet cells in hypoxia treated intestinal organoids (Fig 3B). Paneth cells act as the source of Wnt signaling and secrete Wnt legends. However, our data suggest that this increase in Paneth cells per buds, though, does not compensate for the lower level of Wnt signaling upon hypoxia treatment.

**Figure 3.**
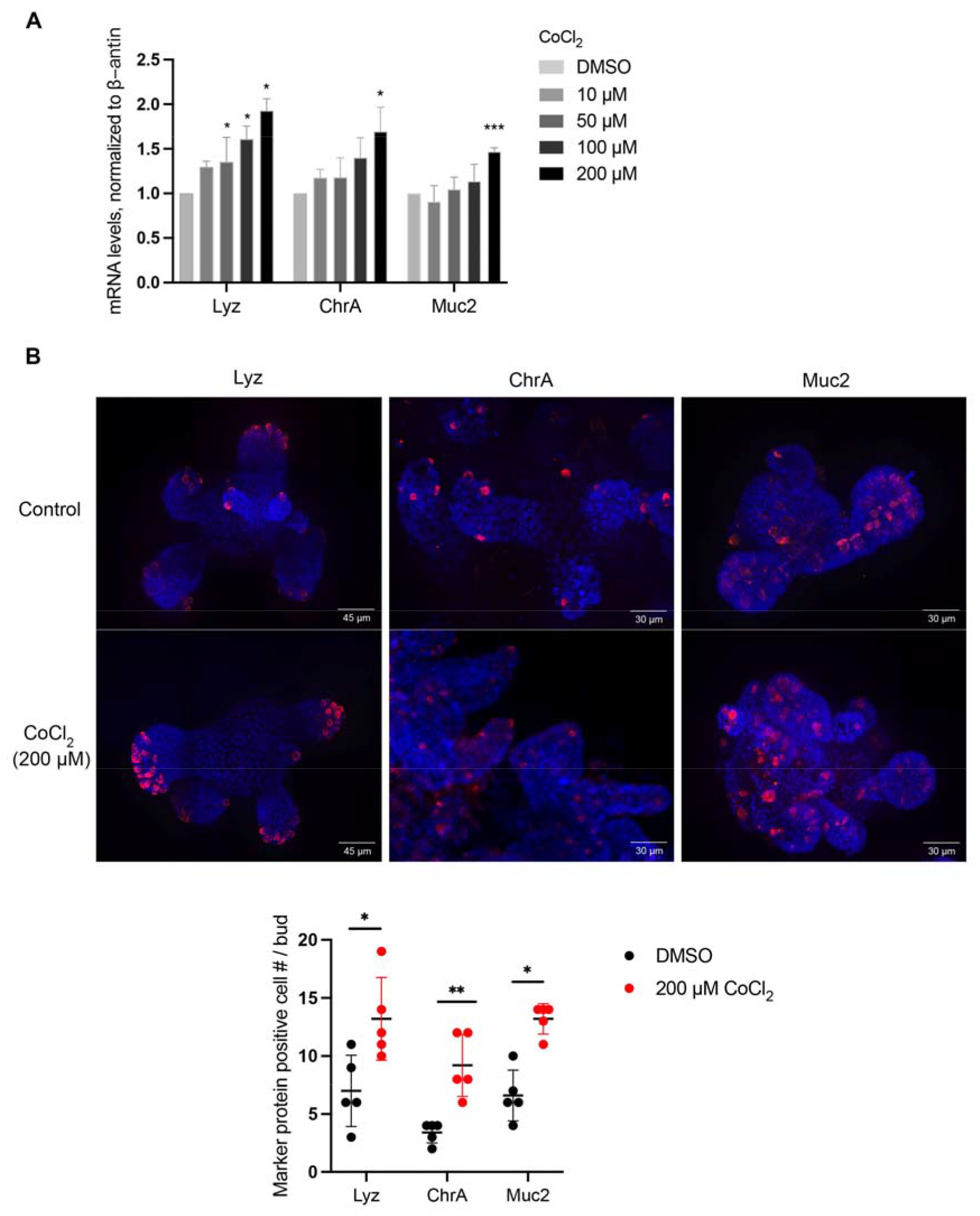
Hypoxia treatment affects intestine differentiation and increases secretary lineages **(A)** RT-qPCR detection of mRNA level normalized to β*-actin* transcript levels of secretary lineage cell marker Lysozyme (Lyz), Chromogranin A (ChrA) and Mucin 2 (Muc2) which marker Paneth cell, enteroendocrine cell and goblet cell, respectively. ^*^*P* < 0.05, ^***^*P* < 0.001. **(B)** Immunofluorescent signal and the quantification by whole mount staining organoids exposed to 200 µm CoCl_2_. Blue indicates DAPI staining, and red indicates signals of Lyz, ChrA and Muc2. Scale bar, 45 μm and 30 µm.

During active inflammation in Crohn’s disease (CD) and ulcerative colitis, pro-inflammatory signals like cytokines promote the stabilization of HIF proteins (45). Hypoxia contributes to inflammation, and Paneth cells are considered to be the origin of intestinal inflammation (46). We then test whether hypoxia induces the inflammatory response in intestinal organoids. Our data showed that hypoxia-induced stabilization of HIF-1α promotes the transcription of proinflammatory cytokines tumor necrosis factor alpha (TNF-α) in intestinal organoids (Fig. 4A, left). This increased TNF-α is likely secreted by the Paneth cells and we wonder if TNF-α would boost this cell lineage. Single intestinal organoid cells treated with a gradient concentration of TNF-α (5-50 ng/ml) for 48 hrs showed increased clonogenic ability, as well as organoid swelling phenotype, especially in the higher TNF-α concentration treatment group (Fig. 4B, C). Interestingly, we notice atypical cell morphology at the bottom of the organoid-Matrigel culture plate in the TNF-α treated cells, which show elongated cell morphology with protrusions (Fig. 4B, red arrows). This boosted growth of organoids probably leads to an adaptation behavior under inflammation assault.

**Figure 4.**
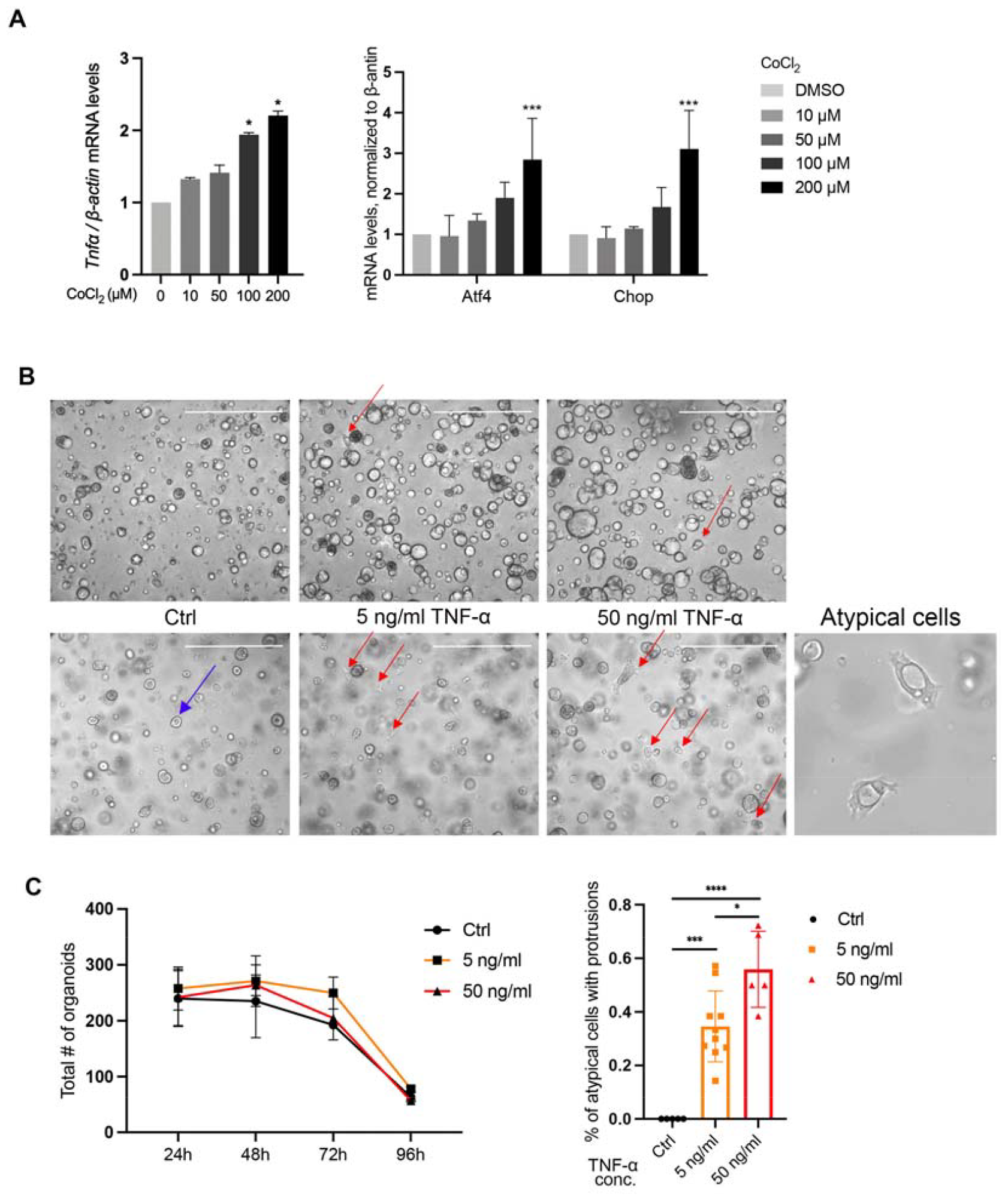
Hypoxia treatment increases the proinflammatory response and UPR response in intestinal organoids **(A)** RT-qPCR detection of mRNA levels of proinflammatory cytokines TNF-α and UPR proteins Alf4 and Chop in hypoxia exposed organoids. ^*^*P* < 0.05, ^***^*P* < 0.001. **(B)** Clonogenicity testing after 5 ng/ml and 50 ng/ml of TNF-α treatment to single organoid cells. Top pictures show the stacked organoids at all layers in Matrigel. The bottom pictures show single layer at the bottom of the culture plate. Blue arrow shows the normal organoid near the bottom of the culturing plate. Red arrow denotes the atypical organoid in TNF-α treated organoids which attach the plate bottom with protrusions. The lower right corner picture shows the representative atypical cells in TNF-α-treated organoids. Scale bar, 1000 µm. **(C)** Left, survival curve showing the clonogenicity of wild type and TNF-α treated organoids for 4 days. Right, statistic of Fig 4B, showing the percentage of the atypical organoids that attach the plate bottom with protrusion.

Paneth cells have active endoplasmic reticulum (ER) functions, and Paneth cell differentiation depends on unfolded protein response (UPR) signaling (47). Unresolved ER stress is common in the IBD epithelial cells (48). Importantly, unresolved ER stress and UPR activation within Paneth cells can be treated as the nidus for intestinal inflammation (46). We detected that mRNA levels of UPR proteins Atf4 and Chop are elevated upon hypoxia exposure (Fig. 4A, right). While X-box binding protein 1 (Xbp1), Grp78 and Perk are not changed significantly (data not shown).

Accordingly, hypoxia treatment increased the secretory lineage cell differentiation in intestinal organoids, further causing the Paneth cell-relative ER stress response.

### 4. Hypoxia treatment reduces the absorptive cell type in intestinal organoids

Enterocytes constitute the most abundant cells and the only absorptive lineage cells in of intestine. In addition to being an element of the epithelial lining, the differentiated mature enterocytes at the upper villi in small intestine play a critical role in digestive, metabolic, and barrier physical integrity maintenance functions (49). They are the cells most susceptible to hypoxia (50). We found that mRNA levels of alkaline phosphatase (Alpi) and fatty acid binding proteins (Fabp1/2) which hallmark enterocytes decreased after the gradient of CoCl_2_ treatment (Fig. 5A). IFC staining also showed a reduced amount of aldolase B (Aldob, marker of enterocyte) positive cells (Fig. 5B), suggesting that enterocyte differentiation is specifically impaired.

**Figure 5.**
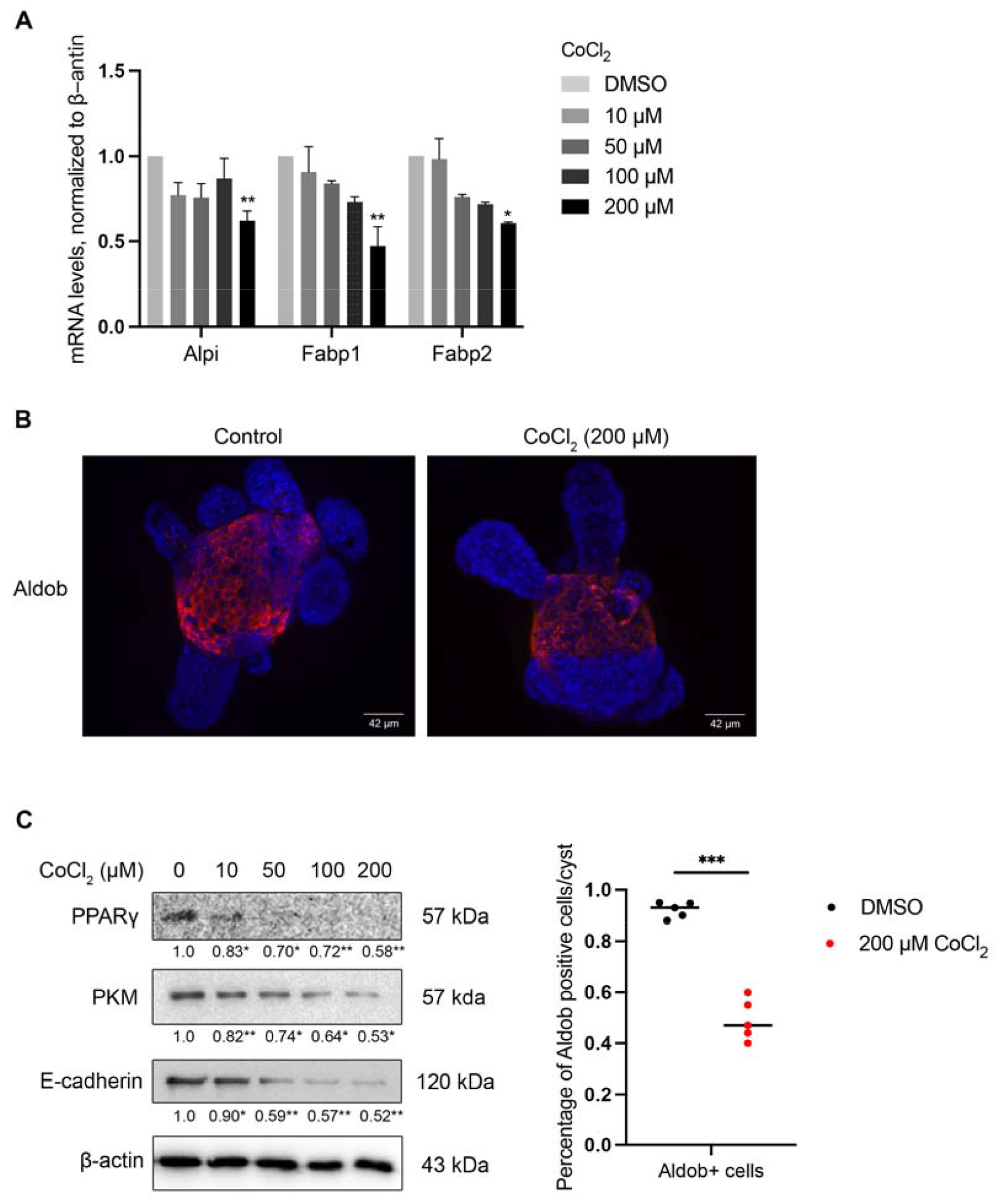
Hypoxia treatment reduces the absorptive cell type in intestinal organoids **(A)** mRNA levels of enterocyte markers of Alpi, Fabp1 and Fabp2 in increasing concentrations of CoCl_2_ treated organoids. ^*^*P* < 0.05, ^**^*P* < 0.01. **(B)** Immunofluorescent signal by whole mount staining of enterocyte marker protein Aldob expression after CoCl_2_ treatment. Blue, DAPI. Scale bar, 42 μm. **(C)** Left, western blot of intestinal barrier relevant protein E-cadherin and metabolism relevant gene PPARγ and PKM in wild-type and CoCl_2_-treated organoids. Numbers below the band indicate the fold change of the band intensity normalized to the β-actin between the control and CoCl_2_ treated groups, which is analyzed by image J. Right, statistic of Fig 5B, showing the percentage of the Aldob positive cells’ area in the cyst structure in the intestinal organoids. ^***^*P* < 0.001.

Enterocytes are involved in nutrient metabolism including glycolysis, fatty acid and lipid metabolism (51, 52). The adipogenic gene Peroxisome proliferator-activated receptor γ (PPARγ) has a beneficial metabolic and anti-inflammatory effect in the GI tract (53). We found that PPARγ, although expressed at a low level in small intestine, was down-regulated after hypoxia treatment (Fig. 5C).

Cells that are exposed to inflammatory cytokines manifest increased glycolysis, which is reminiscent of those observed in hypoxic cells (54, 55). Glucose is also a relevant energy source for enterocytes, and pyruvate kinases are responsible for catalyzing the final step in glycolysis (56). We detected the protein level of glycolysis gene pyruvate kinases M (PKM) was reduced after hypoxia exposure (Fig. 5C).

In addition to the general structure of intestinal organoids as the barrier, which we detected no change after hypoxia treatment (Fig. 1A), we wondered whether the intrinsic elements of the epithelium barrier were affected. We detected hypoxia caused a gradient down-regulation of the epithelial barrier protein E-cadherin (Fig. 5C), which might indicate higher permeability in the gastrointestinal epithelium

Therefore, hypoxia exposure decreased the absorptive enterocytes, with reduced metabolic-related proteins and might dampened intestinal barrier function.

## Discussion

Hypoxia occurs in intestinal diseases such as IBD and NEC (22, 23). It is well established that hypoxia-induced intestinal injuries and low oxygen-relative physiological responses (23). Studies using murine model of colitis indicated that HIF-1α plays a protective role in intestinal inflammatory mucosal disease, most of which suggested a barrier-protective function of HIF-1α (16, 20, 22). As different intestinal lineage cells executive specific functions, however, the mechanism of hypoxia caused various cellular or metabolic responses failed to be elucidated at the cell-type specific level in IEC. In this study, we showed that hypoxia exposure by hydroxylase inhibitor CoCl_2_ significantly changes the small intestinal organoid cell lineage differentiation and reduces the ISC function by down-regulating Wnt signaling.

It’s known that hypoxia contributes to colorectal cancer cell (CRC) and activates Wnt signaling (57, 58). However, in normal intestine epithelium under homeostasis, it’s less clarified whether hypoxia impacts ISC function and alters Wnt signaling. Consistent with Liu and Zu et al. showing that Wnt signaling is reduced in the hypoxia-induced NEC and ischemia-reperfusion (I/R) murine model (59, 60), our results further demonstrate that hypoxia-caused impaired ISC function was accompanied by reduced Wnt signaling.

This intestinal organoid-based hypoxia exposure model provides an opportunity with the cell type-specific model which correlates HIF-1 and the target genes in inflammation or metabolism dysfunctions.

Notch signaling plays a crucial role in regulating cell fate decisions in intestinal homeostasis (61, 62). Upon binding to their ligands, Jagged (Jag) and Delta-like (Dll) proteins, the Notch receptor was cleaved and the intracellular domain translocates to the nucleus, where it activates the transcription of the target genes, such as hairy and enhancer split-1 (Hes1), regulating enterocyte differentiation (63). In reverse, Hes1 represses the expression of atonal homolog 1 (Atoh1), which regulates secretory cell lineage differentiation.

Studies have implied that hypoxia increases the expression of Notch direct downstream genes in neuronal and myogenic cells (64). Enterocyte differentiation is impaired as a consequence of reduced Notch signaling pathway in patients with CD which is accompanied by hypoxia-featured inflammation (65). While Notch1 signaling is upregulated in colorectal hypoxic Caco-2 cells (66). It is still controversial regarding the role of Notch signaling in regulating hypoxia-relevant intestinal diseases. Yeung et al. reported that hypoxia increases clonogenicity and maintains the stem-like phenotype of colorectal cell line-derived cancer stem cells, preventing differentiation of enterocytes and goblet cells by activating Notch1 (67). Of note, it remains less reported about the relationship between hypoxia and Notch signaling in mediating stemness or barrier function in intestine under homeostasis. Although we found the absorptive enterocytes decrease after hypoxia treatment, its regulatory Notch signal transcription factors such as Hes1 and Notch intracellular domain (NICD) should be tested in the future to answer if hypoxia affects the Notch signal directly.

Cell cultures have been used to study intestinal epithelial permeability after repair/restitution (68, 69), while intestinal organoids provide us better *in vitro* model to study the mechanism of injuries such as hypoxia on intestinal epithelial barrier function which can best mimic the *in vivo* intestine. In intestinal cell lines or *in vivo* hypoxia-exposed models (70, 71), HIF-1α was reported to influence intestinal epithelial permeability (19, 72). Liu et al. suggested that Notch1 signaling plays a role in maintaining the intestinal barrier function by regulating TJ proteins (66). More experiments are expected in the future such as testing the expression of TJs (Zo-1 and occludin) and intestinal mucosal permeability alterations, to prove the regulation of Notch to TJ proteins and its role in maintaining barrier function.

On the other hand, we detected a higher number of goblet cells with increasing Muc2 level after hypoxia exposure, consistent with D. Ortiz-Masiá et al.’s finding with increased Muc2 in chronic CD patients (65). Goblet cells secret the mucins that form the physical epithelial barrier, and Paneth cells secrete antimicrobial and some signaling factors to regulate the ISC stemness (73). The differentiation of IECs from the ISCs into the terminal differentiated cells is under the regulation of master transcription factors (74). The progenitor cells committed to secretory lineage have high expression of Atoh1. GFI1 and Spdef specify goblet and Paneth cells, Neurogenin3 and NeuroD1 mark enteroendocrine cell differentiation, and Elf3 and KIf4 regulate goblet cell differentiation. More investigation is expected to delineate whether hypoxia shifts early differentiation toward the secretory lineage via changing the upstream secretory transcription factors.

Cells undergo stress and increase the production of certain proteins in their endoplasmic reticulum (ER) when they experience stressful conditions like hypoxia (75), which is known as the unfolded protein response (UPR). Heijmans et al. showed ER stress plays an important role in regulating normal intestinal epithelial stem cell differentiation (47). We detected the UPR targets Chop and Atf4 are up-regulated after hypoxia exposure. As XBP1 is a regulator of the unfolded protein response, it’s required for ER expansion (Shaffer et al., 2004) and adaptation of tumor cells to hypoxic conditions (48). XBP1 deletion induces inflammation in intestine (48). Although XBP1 was reported to drive the triple-negative breast cancer (TNBC) tumorigenicity by controlling the HIF1α pathway (76), however, we observed the level of Xbp1 did not change obviously in hypoxia-exposes normal intestinal organoids.

As hypoxia treatment dampens intestinal organoid clonogenicity, while TNF-α treatment boosts it, they likely target different cell types in the intestinal organoids. Compared with the absorptive cell types (i.e., enterocytes) as the most vulnerable cells under hypoxia threat, Paneth cells being the major warriors in front of TNF-α might explain the opposite growth trend under these two different insults. Because TNF-α treatment 2 days post single organoid seeding has the opposite barrier function protein alteration compared with those single-cell formed organoids (data not shown), further enhancing the possibility that TNF-α might boost Paneth cells in the first two days of organoid growth (wherein stem cells are the majority cells) while mainly kill the enterocyte cell at later days. These two different insults targeting different intestinal cell types help to explain that each cell type performs its function under homeostasis and pathologic conditions. Besides, as we also notice atypical cell protrusions in TNF-α treated cells, further experiments such as hypoxia and TNF-α co-treatment can be done to elucidate whether this inflammation-fueled cell type-specific outcome helps intestinal cancer cell metastasis, and whether hypoxia enhanced the secretive cell number can further promote this phenomenon.

## Method

### Animals and crypt isolation organoid culture

Small intestines were removed from sacrificed 4-16-week-old C57BL/6 mice, flushed in cold PBS and cut open. Mesentery was removed and the tissue was washed three times before being cut into pieces. Then the tissues were digested with 2 mM EDTA and stayed on ice for 45 min. The digested intestines were transferred into a tube with cold PBS and shaken around 3 times to detach the villi and remove residual EDTA. The suspension was filtered through 70-μm meshes (BD Falcon) to enrich crypts. The crypt-containing solution was centrifuged at 150 g for 3 min. The crypt pellet was then resuspended with DMEM and embedded in Matrigel (Corning, #356231) for solidification for 15 min in a 37° C incubator. Then culture medium was added above the crypt-Matrigel mix with DMEM/F12 (Gibco) medium supplemented with EGF (50 ng/ml, Invitrogen), Noggin (50 ng/ml, R&D SYSTEMS), R-spondin 500 ng/ml (R&D SYSTEMS), 1 × N-2 (Invitrogen), 1 × B-27 (Invitrogen), HEPES (10 mM), 1 x GlutaMax (Gibco), penicillin and streptomycin (100 U/ml), N-Acetyl Cysteine (1 μM, Sigma). Y27632 (10 μM, Selleck) was included in the culture medium during the first 2 days after being passaged. The medium was changed every other day.

### Hypoxia treatment

Organoids were treated with gradient concentrations of CoCl_2_ (Sigma, #V900059) by 0, 10, 50, 100, and 200 μM for 24 hours. DMSO (concentration less than 0.1%) was used as the solvent and control reagent.

### Clonogenicity assay

The colony-forming experiment was measured by seeding 200–300 single-cell organoids in Matrigel and calculating organoid number after 2 days. The organoids were cultured for five to nine days, then the crypt number was collected each day. Single organoid cells were obtained by TrypLE (ThermoFisher) treatment for 10 min at 37°C, followed by gentle pipetting if cell clusters remained.

### RNA Isolation and Reverse transcription-quantitative polymerase chain reaction (RT-qPCR)

Total cellular RNA was extracted using EZ-10 RNA Mini-Preps Kits Handbook, (#BS88583, Markham ON, Canada). Around 250 ng RNA was used to do reverse transcription by RevertAid First Strand cDNA Synthesis Kit (K1622, Thermo, Waltham, MA, USA). Primers are shown in Table 1. RT-qPCR was performed using an SYBR qPCR Master Mix (Q321-03, Vazyme, Nanjing, China) by the following conditions: pre-heat at 95°C for 10 s, then followed with 39 cycles at 95°C for 5/10 s and 60°C for 30 s (72°C for 30 sec). The relative mRNA levels were analyzed using the 2^−ΔΔCt^ method. Experiments were repeated at least three times. Then the mean value was used to calculate the significant alterations in statistical analysis.

**Table 1.**
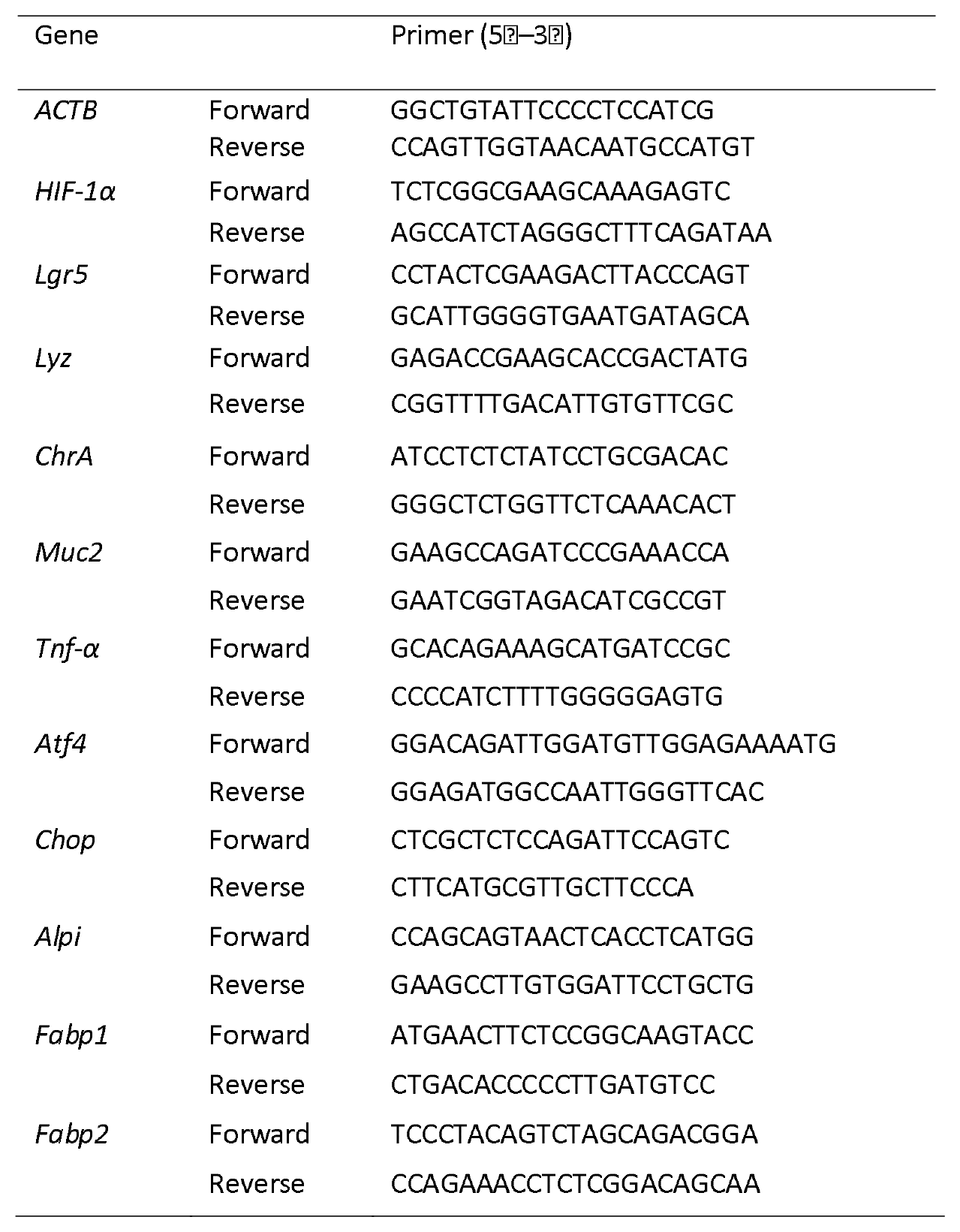
Primers used for RT-qPCR.

### Whole mount staining and Immunofluorescence analysis

Organoids were cultured in 6-well plate containing 2 ml medium. Four days after passaging and then treated with CoCl_2_ for 24h, organoids were collected in ice-cold culture medium and Cell Recovery Solution (BD Biosciences) was added and incubated for 1 hour to remove the Matrigel. After being washed 3 times by medium, the organoids were fixed by 2% formaldehyde overnight at 4°C. On the second day, organoids were permeabilized in 0.1% Tween 20/PBS for 30 min at room temperature. Blocking was done by 0.2% Bovine serum albumin (BSA) for one hour at 4°C. Then organoids were incubated in 1st antibody at 4°C overnight. On the third day, three times of washing were performed with ice-cold 0.1% Tween 20/PBS, followed by organoid sedimentation at room temperature. Then secondary rabbit or mouse anti-rat IgG (H+L) reagent was added to incubate for 1 hour, followed by three times washing with 0.1% Triton X-100. Finally, the pelleted organoids were mounted on the glass slides. Images were taken by Spinning Disk Confocal Microscopy and five random fields were chosen to observe the 1st antibody (Lyz, ChrA, Muc2, and Aldob) positive cells and measure their percentage. The experiment was repeated for at least three times.

### Western blotting

Organoids cultured in the 24-well plate were washed by cold PBS and moved into 1.5 ml tubes. Supernatant with Matrigel was removed and 80 µl 4X loading buffer (with DTT) was added and pipetted for mixing. Samples were denatured at 100°C for 10 min. Protein samples were stored at -80°C if not used immediately. Protein samples (around 50 µg) were loaded and for electrophoresis on 8% sodium dodecyl sulfate-polyacrylamide gels, followed by transferring to a polyvinylidene fluoride (PVDF) membrane (Bio-Rad, CA, USA). Then the blocking was carried in non-fat dry milk at 37°C for 1 h. First antibody incubation was done at 4 degrees with the following antibodies (1:1000). The rabbit monoclonal anti-HIF-1α (#36169T), anti-β-Catenin (#8480) and anti-Cyclin-D1 (#2978) were purchased from Cell Signaling Technology (Devers, MA, USA). The rabbit monoclonal anti-Axin2 (#2151) was obtained from Abcam (Cambridge, UK). The mouse monoclonal anti-E-cadherin was purchased from BD Biosciences (610181, San Jose, CA, USA). The mouse monoclonal, anti-Ephb2 (sc-130752), anti-PKM (sc-365684) and anti-PPARγ (sc-7273) were purchased from Santa Cruz (Dallas, TX, US). After 1^st^ antibody incubation, membranes were washed by Tris-Buffered Saline Tween (TBST)-20 for 3 times. Incubation with the secondary goat anti-rabbit or goat anti-mouse IgG (1:1000, ZSGB-BIO, Beijing, China) were performed at 37°C for 1 h. Then the PVDF membranes were washed three times for 10 min each time. The blots were developed using an enhanced chemiluminescence detection kit (Millipore, MA, USA). Image J was used to do the densitometric analysis.

### Statistical analysis

GraphPad Prism 9.0 was used to perform statistical analysis. one-way ANOVA was used to test the differences among groups, which is followed by Tukey’s multiple comparisons test. Each experiment was repeated at least 3 times. The results were deemed significantly different at P < 0.05.

## Author contributions

Xi Lan designed the paper, Xi Lan, Qiu Ping and Mou Chunfeng did the experiments, and Mou Chunfeng approved the version to be published. All authors approved this version for publication.

## Acknowledgment

May this tiny work find the sincerest passion and effort of those unrooted scientists, in honor of their unseen sweat and industrious missing contribution.

